# PHACTboost: A Phylogeny-aware Boosting Algorithm to Compute the Pathogenicity of Missense Mutations

**DOI:** 10.1101/2024.01.30.577938

**Authors:** Onur Dereli, Nurdan Kuru, Emrah Akkoyun, Aylin Bircan, Oznur Tastan, Ogün Adebali

**Affiliations:** Faculty of Engineering and Natural Sciences, Sabanci University, Istanbul 34956, Turkiye; TUBITAK-ULAKBIM Turkish Academic Network and Information Center, Ankara 06530, Turkiye; TUBITAK Research Institute for Fundamental Sciences, Gebze 41470, Turkiye

**Author notes:** These authors contributed equally to this work. Co-first authors O.D. and N.K. have the right to list themselves first in author order on their CVs.

**Keywords:** phylogenetics, Mendelian diseases, pathogenicity scoring, amino acid substitution, machine learning, gradient boosting

## Abstract

Most algorithms that are used to predict the effects of variants rely on evolutionary conservation. However, a majority of such techniques compute evolutionary conservation by solely using the alignment of multiple sequences while overlooking the evolutionary context of substitution events. We had introduced PHACT, a scoring-based pathogenicity predictor for missense mutations that can leverage phylogenetic trees, in our previous study. By building on this foundation, we now propose PHACTboost, a gradient boosting tree-based classifier that combines PHACT scores with information from multiple sequence alignments, phylogenetic trees, and ancestral reconstruction. The results of comprehensive experiments on carefully constructed sets of variants demonstrated that PHACTboost can outperform 40 prevalent pathogenicity predictors reported in the dbNSFP, including conventional tools, meta-predictors, and deep learning-based approaches as well as state-of-the-art tools, AlphaMissense, EVE, and CPT-1. The superiority of PHACTboost over these methods was particularly evident in case of hard variants for which different pathogenicity predictors offered conflicting results. We provide predictions of 219 million missense variants over 20,191 proteins. PHACTboost can improve our understanding of genetic diseases and facilitate more accurate diagnoses.

---

Missense mutations can alter protein function, activity, and stability by modifying the amino acid sequence in the coding regions (Stefl et al., 2013; Thusberg & Vihinen, 2009). Many missense variants are linked to diseases (Stefl et al., 2013; Wang & Moult, 2001), and better understanding their potential to cause diseases (pathogenicity) is important. However, the state of pathogenicity of a vast number of variants remains undetermined at present. Computational tools can help predict the pathogenicity of these variants (Carter et al., 2013; Eilbeck et al., 2017), and many computational approaches have been developed to this end (Adzhubei et al., 2013; Calabrese et al., 2009; Ioannidis et al., 2016; Jagota et al., 2023; Malhis et al., 2020; Ng & Henikoff, 2003; Pejaver et al., 2020; Raimondi et al., 2017; Sim et al., 2012; Wu et al., 2021).

Currently available tools for pathogenicity prediction can be broadly categorized into two groups: methods based on conventional statistical approaches (Davydov et al., 2010; Malhis et al., 2020; Ng & Henikoff, 2003), and machine learning-based methods (Adzhubei et al., 2013; Carter et al., 2013; Ioannidis et al., 2016). Some of these machine learning tools include the output of other learning-based pathogenicity predictors into their training model to enhance the prediction ability (namely, meta-predictors) (Ioannidis et al., 2016). Many traditional statistical methods for predicting pathogenicity depend on the degree of conservation of protein position derived from multiple sequence alignments (MSAs). The integration of features based on evolutionary conservation along with structural, functional, and physicochemical attributes into an algorithm has the potential to enhance the capabilities of tools for predicting pathogenicity. This insight has led to the introduction of machine learning-based approaches to the domain.

The machine learning-based approaches used in the area can be divided into two groups: supervised methods, which are primarily trained by using human variants, and unsupervised methods, which encompass such techniques as autoencoders and language models. We apply a diverse set of tools sourced from the dbNSFP (Liu et al., 2020), a well-established database containing the results of tools to predict the effects of variants, in a comparative analysis in this study. This set includes tools reported in the dbNSFP as well as novel techniques, such as AlphaMissense (Cheng et al., 2023), EVE (Frazer et al., 2021), and CPT-1 (Jagota et al., 2023). The latter set of tools are known for their exceptional performance in predicting pathogenicity compared with prevalent methods in the area. AlphaMissense is a recently developed method that uses unsupervised protein language modeling and the system for predicting the structure of proteins used in AlphaFold (Jumper et al., 2021). On the contrary, EVE is an unsupervised deep generative model, while CPT-1 provides a novel approach based on cross-protein transfer learning by leveraging deep mutational scanning experiments (DMS) data from five proteins. We include these tools in our comparative analysis owing to their impressive performance and the significant impact that they have had on research in the field.

Evolutionary conservation is a major information source used by many predictive tools. However, these methods rely solely on MSAs instead of the phylogenetic tree when computing evolutionary conservation, and thus, fall short of accurately predicting conservation at the given position. An illustrative example is given in Fig. 1A. When phylogenetic trees are not considered, the two scenarios shown in the figure are equivalent in terms of the observed frequency of histidines (Hs) and glutamic acids (Es), representing the reference and the alternating amino acids, respectively. On the contrary, the two cases are entirely different when the context of the change in phylogenetic trees are taken into account. The alternating amino acid E is observed at four species (the leaves) in the first tree independently. We refer to these types of alterations as “independent events.” While substitution to E is also observed four times in the second tree, all instances of it is descendant of a single mutation in the same ancestor, which we refer to as a “dependent event.” When the evolutionary context is not considered, the H to E alteration may appear equally possible. However, when the evolutionary history considered, there is more evidence that this change would be neutral in the first tree as this changed occurred repeatedly and independently. Additionally, as in the first example in Fig. 1A, observing the alternative amino acid in a close species to human on the corresponding phylogenetic tree, provides more evidence on the neutrality of the change. We recently introduced PHACT, a phylogeny-based algorithm to predict the effects of variants (Kuru et al., 2022), to overcome the limitations of prevalent tools that ignore independent and dependent alterations. PHACT scores the amino acid substitutions based on phylogenetic trees and the probabilities of the ancestral reconstruction of amino acids. It considers the reconstructed distribution of ancestral amino acids in each species and their distance to the human leaf. This allows for an assessment of the nature of change in the amino acid as well as the relevance of this change to the human protein. We demonstrated that PHACT outperforms conventional statistical methods provided in the dbNSFP.

**Figure 1.**
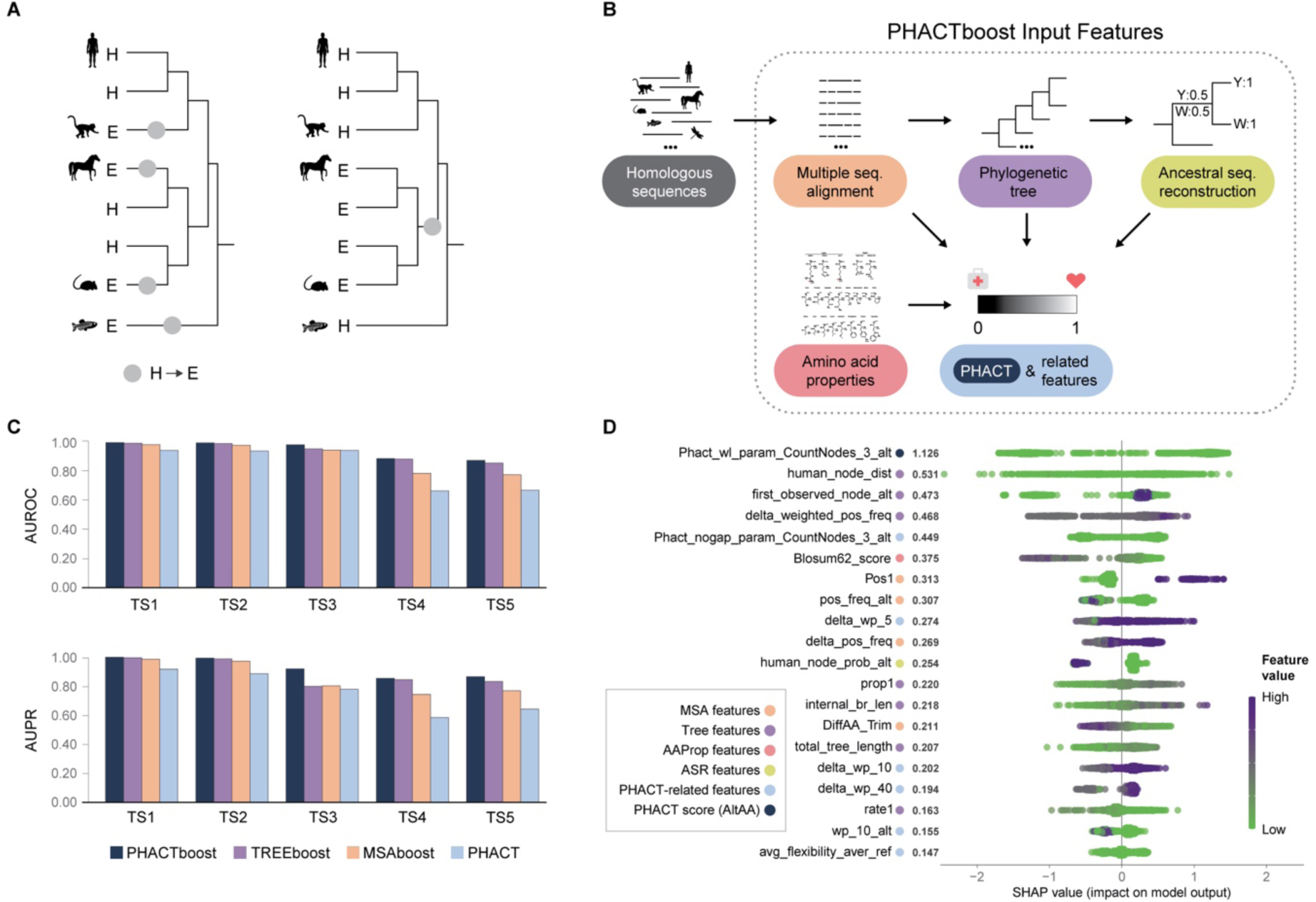
**A.** Motivation for designing a phylogeny-based approach. The green circles represent the locations of alterations in amino acids from H to E. **B.** Outline of the proposed PHACTboost algorithm. The main input to the model consists of PHACT and PHACT-related scores. We provide other features derived from the MSA, phylogenetic trees, and ancestral sequence reconstruction (ASR). Because we limit our BLAST search to a maximum of 1,000 hits, there are cases for which we do not obtain the full evolutionary history. The properties of the amino acid serve as an additional type of feature in such cases. **C.** Evaluation of the performance of the PHACTboost algorithm encompassing all alternative test sets. It reveals the improvements attained over the baseline PHACT model, MSAboost, and TREEboost models that apply the same machine learning algorithm but differ in their feature sets. MSAboost uses only MSA-based features while TREEboost uses both MSA- and tree-based features. The AUROC and AUPR levels of all three tools are given in Supplementary Table 1. **D.** Exploration of feature importance– Shapley values of the PHACTboost algorithm. The features have been categorized into five distinct groups, represented by colors similar to those in Figure 1B. More information about the input features and their respective groups can be found in Supplementary Table 2.

By building on the foundation of incorporating the evolutionary context into predictions of the effects of variants (Kuru et al., 2022), we propose PHACTboost, an improved version of PHACT, in this study. Because PHACT is score based, it does not learn from the large set of available data on pathogenic and neutral variants. PHACTboost integrates PHACT scores, the components of PHACT, and other information computed on phylogenetic trees, MSA, and ancestral probability distributions in a machine learning framework. Figure 1B illustrates the input files used for feature construction in PHACTboost, with five distinct colors representing different sources of information. PHACTboost uses a gradient boosting-based model of learning that is trained on various sets of features computed over the input sequence and mutations. We also thoroughly assess the predictive performance of PHACTboost. The results showed that it can outperform 40 algorithms to predict pathogenicity reported in the dbNSFP (version 4.4a; whenever we refer to the dbNSFP, we refer to version 4.4a unless stated otherwise).

We also compared PHACTboost with more recently developed tools to predict pathogenicity that are not part of the dbNSFP, including AlphaMissense, EVE, and CPT-1. The results showed that PHACTboost delivered superior predictive performance across AUC and AUPR levels on different subsets of variants. We also report the specific improvement in performance brought about by it over established methods reported in the dbNSFP as well as the above-mentioned recently developed tools. We conducted experiments on various subsets of test data to verify the validity of this conclusion. The improvement in performance due to PHACTboost was more pronounced on hard instances that yielded the most conflicting results among the relevant tools. Furthermore, the pathogenicity scores of PHACTboost were correlated with DMS experiments, and delivered an average Spearman correlation comparable to those of the highly successful tools AlphaMissense, CPT-1, EVE, and REVEL (Ioannidis et al., 2016). These findings underscore the effectiveness of PHACTboost in accurately predicting the pathogenicity of missense mutations and highlight its potential as a valuable tool in variant prioritization for the diagnosis of genetic diseases.

We provide details of our contributions in the Results section below. We first elaborate on the development and features of PHACTboost. Subsequently, we discuss the roles of various features in pathogenicity prediction. Following this, we present the results of our comparative analysis, and showcase how PHACTboost outperformed other tools reported in dbNSFP as well as such novel tools as EVE, CPT-1, and AlphaMissense. Finally, we analyze variants that were misclassified by PHACTboost.

## Results

We conducted extensive computational experiments to assess the predictive performance of PHACTboost. We trained it on a dataset of 70,937 variants from 13,062 proteins, including 31,942 neutral and 38,995 pathogenic variants. The main test set (referred to as TS1) comprised 14,116 neutral and 13,194 pathogenic variants belonging to 8,423 proteins. The dataset of variants included a set of clinically verified pathogenic variants obtained from ClinVar, and neutral variants with an allele frequency (AF) higher than 0.01 from ClinVar (Landrum et al. 2016) and gnomAD (Karczewski et al. 2020). The construction of the training and test sets is detailed in the “Methods” section.

We also established subsets of increasing difficulty within TS1 to further assess the capabilities of generalization of the models. To avoid circularity caused by the presence of the same proteins both in the training and the test sets, we created TS2 and TS3 that comprised variants of proteins and variants located on protein positions that were not present in the training dataset, respectively. We also created TS4 and TS5 to assess the performance of PHACTboost and other tools on the hardest cases. TS4 included variants, on the predictive classification of which at least 25% of the 40 tools in the dbNSFP disagreed (i.e., whether they were pathogenic or neutral). TS5 focused on mixed proteins that contained pathogenic and neutral variants from the TS4 dataset and yielded the most challenging variants to predict. Table 1 provides a comprehensive breakdown of the compositions of the training set and the alternative test set. It is important to note that the same training set was used for all evaluations in this study.

**Table 1.** Summary of the datasets constructed to evaluate the discriminative capabilities of PHACTboost and benchmark algorithms.

| Sub-Dataset | Number of Proteins | Number of Pathogenic Variants | Number of Neutral Variants | Dataset Explanation |
| --- | --- | --- | --- | --- |
| Training Set | 13,062 | 38,995 | 31,942 | The main sets are ClinVar and gnomAD. HSV is used to exclude variations with conflicting interpretations in the literature. |
| TS1 | 8,423 | 13,194 | 14,116 | The main test set including pathogenic variants added to ClinVar after 2020. |
| TS2 | 8,302 | 9,830 | 13,913 | Variants located on protein positions not observed in the training data |
| TS3 | 913 | 264 | 1,291 | Variants from proteins that are not present in the training data |
| TS4 | 923 | 366 | 696 | Variables in which at least 25% of dbNSFP tools predict the class labels discordant. |
| TS5 | 561 | 254 | 393 | Composed of mixed proteins included in TS4 that exhibit both neutral and pathogenic variants |

We used the ROC and the PR curve as well as the corresponding values of the AUROC and AUPR to assess the performance of the methods in terms of predicting pathogenicity. All performance measures are reported on this set unless otherwise stated. Moreover, we focused on comparing the performance of PHACTboost with that of the state-of-the-art tools AlphaMissense, EVE, and CPT-1.

### PHACTboost provides an improvement over PHACT

We first compared the performance of PHACTboost with that of PHACT, as shown in Fig. 1C. PHACTboost achieved values of the AUROC and AUPR of 0.99, while PHACT recorded an AUROC of 0.937 and an AUPR of 0.909 on TS1. The significant gain in performance achieved by PHACTboost over PHACT was consistently observed across all subsets of the test set (Fig. 1C).

To further examine the contributions of features of ancestral reconstruction and PHACT-related features, we compared the performance of PHACTboost with that of a model that relied only on MSA-based features. We refer to this model as “MSAboost,” and Fig. 1C shows that PHACTboost achieved an AUROC that was 1.4% higher than that of MSAboost on TS1. We observed a similar trend of the disparity in performance between PHACTboost and MSAboost across TS2 and TS3, with differences of 1.6% and 3.7%, respectively. These results indicate that while the classifier could successfully learn by using MSA-related features, the additional features of ancestral reconstruction and PHACT-based features enabled PHACTboost to deliver better performance. This gain in performance was more pronounced on TS4 and TS5. Fig. 1C shows that PHACTboost achieved AUROC scores that were 10.1% and 9.7% higher, and AUPR scores that were 11.1% and 9.7% higher than those of MSAboost on TS4 and TS5, respectively. These two datasets are particularly important because guidelines of the American College of Medical Genetics (ACMG) regard the outcomes of established computational tools on them as evidence of their accuracy in terms of identifying the types of variants. Most computational tools tend to produce similar predictions as a result of this influence. This similarity arises because their predictions have already been considered when assigning labels to the variants submitted to ClinVar. We constructed the datasets TS4 and TS5, which we call sets of “hard cases,” by using a set of variants that were free from this effect as much as possible.

We designed TREEboost, a model based on both the MSA and tree-based features, as a second alternative approach to highlight the importance of PHACT-related features and features of ancestral reconstruction. We observed differences of 0.4%, 0.4%, 2.9%, 0.3%, and 1.7% between the AUROC scores of PHACTboost and TREEboost on TS1, TS2, TS3, TS4, and TS5, respectively. The gain in performance due to PHACTboost was clearer in terms of the AUPR levels. Fig. 1C and Supplementary Table 1 show that the corresponding differences in the AUPR between PHACTboost and TREEboost on TS1–TS5 were 0.4%, 0.6%, 11.9%, 1%, and 3.3%, respectively.

### Contributions of features to pathogenicity prediction

Fig. 1D presents Shapley values to provide insights into the contribution of each feature to the predictive performance of the proposed tool. The 20 features with the most significant influence on the inputs to the model are shown in the figure. The PHACT score stood out as the most influential feature in terms of pathogenicity prediction, and this underscores its crucial role in the performance of the model. To gauge the value of each source of information, we categorized the features into five distinct groups, represented by colors similar to those in Figure 1B. The details of the input features and their corresponding classes are given in Supplementary Table 2. Notably, 15 of the 20 most influential features were related to tree- and ancestral reconstruction-based features. This shows the significance of incorporating phylogenetic trees and evolutionary information into the model.

### PHACTboost outperforms approaches reported in dbNSFP

Fig. 2 and Supplementary Fig. 1 present one-to-one comparisons of PHACTboost with 40 tools reported in the dbNSFP, with the protein majority vote (PMV) as the predictor in terms of the AUROC on various subsets of the test set. Please note that for the tools that have multiple versions, we included the one with the highest performance in the figure, yielding 27 unique tools. Not all tools provided predictions for the entire set of the test dataset of variants, and the test set became too small when we intersected the set of variants to create a common set. To maximize the portion of the test data evaluated by each tool, we created subsets of our test set for each one-to-one comparison based on variants commonly reported by the compared pair of tools. Note that this resulted in different portions of the test data. We quantified the differences in performance through ΔAUROC (ΔAUPR) levels—the difference between the AUROC (AUPR) of the corresponding tool and that of PHACTboost. Negative values showed that the PHACTboost algorithm exhibited superior performance. The resulting ΔAUROC levels are shown in Fig. 2, while Supplementary Fig. 1 displays the ΔAUPR levels. Further details related to these comparisons, such as the AUROC and AUPR of each tool, as well as the number of neutral and pathogenic variants in each comparison are listed in Supplementary Tables 3 and 4 for each subset generated.

**Figure 2.**
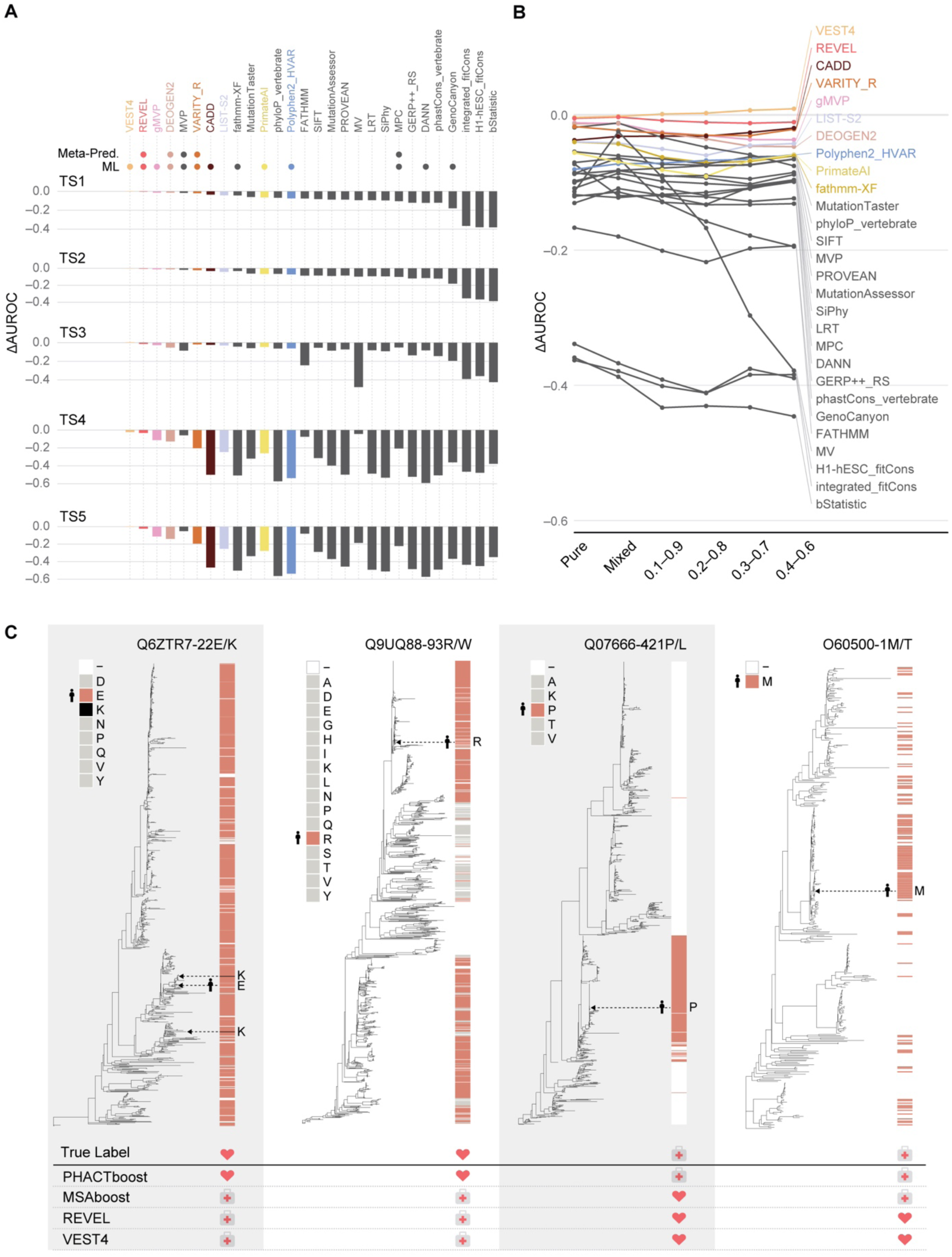
AUROC comparisons of PHACTboost with tools reported in the dbNSFP and PMV. **A.** Comparisons on TS1, TS2, TS3, TS4, and TS5, respectively. ΔAUROC corresponds to the difference between the AUROC levels of the other tools and PHACTboost, and negative values represent better predictive performance by PHACTboost. Each tool is annotated with respect to whether it is a meta-predictor, based on machine learning, or a traditional statistical approach (see Supplementary Table 5). **B.** Comparisons on TS1, TS1_Pure, TS1_Mixed, TS1_0.1-0.9, TS1_0.2-0.8, TS1_0.3-0.7, and TS1_0.4-0.6. **C.** Example of variants misclassified by VEST4 and REVEL but correctly predicted by PHACTboost. The original labels and those predicted by the tools are depicted in gray for pathogenic variants and coral for neutral variants. The colors corresponding to the amino acids in the MSA are displayed next to each subplot. Coral represents the reference while black denotes alternating amino acids. The human leaf is marked on each tree, and the UniProt ID, position, and reference of the protein as well as alternative amino acids are annotated next to each plot.

Fig. 2A shows that PHACTboost outperformed the benchmark tools on all subsets except TS3. Only VEST4 (Carter et al., 2013) recorded slightly higher (0.004%) AUROC levels than PHACTboost on TS3. However, the difference between them was minimal, and the training set of VEST4 was not accessible. Consequently, it is possible that the test data might have overlapped with its training set to some extent.

Grimm et al. (Grimm et al., 2015) showed the impact of the composition of a given dataset on the performance of predictive methods. Proteins containing predominantly neutral or pathogenic variants in the training set in particular yield optimistic predictive results on test variants belonging to them. Our earlier experiment on “unseen proteins” (TS3) is one way to factor out such easy cases. Furthermore, we evaluated the performance of our algorithm on various subsets generated based on the PMV. The details of the subsets are given in the “Methods” section. The PMV is the ratio of pathogenic-to-neutral variants within a protein’s variants observed in the training set. A larger fraction of proteins with a more balanced pathogenic-to-neutral ratio renders the dataset more reliable for performance comparisons. Fig. 2B and Supplementary Fig. 1B show the gain in the performance of PHACTboost against tools reported in the dbNSFP and PMV on subsets generated by using different ranges of the PMV. VEST4 recorded slightly better AUROC levels on the subsets, with pathogenic-to-neutral ratios of 0.2 to 0.8, 0.3 to 0.7, and 0.4 to 0.6. However, as mentioned earlier, the lack of access to the training set of VEST4 might have led to its optimistic evaluation. Overall, the superior performance of PHACTboost compared with the other tools support our claim that incorporating evolutionary information and considering independent substitution events, achieved by using PHACT scores and PHACT-related features, improves the accuracy of predictions of the effects of variants.

We observed that most prevalent tools tended to mislabel i) neutral variants that were observed in a species close to human, but were rare in terms of frequency, and ii) pathogenic variants that were detected in multiple species because of a single mutation in their ancestral species. We further investigated some neutral and pathogenic variants that had been mislabeled by MSAboost, VEST4, and REVEL, which delivered exhibited superior performance to tools reported in the dbNSFP as shown in Fig. 2 but were correctly classified by the PHACTboost and PHACT algorithms (Fig. 2C). We detail these cases below.

The first two cases involved samples of variations labeled as pathogenic by VEST4, REVEL, and MSAboost, but reported as neutral in ClinVar. The first case corresponded to a substitution from E to K at position 22 of the protein with UniProt ID Q6ZTR7. Although K was observed only twice, it appeared at leaves close to the human leaf and twice appeared independently. Each piece of information highlights the neutrality of the variation. The second case consisted of an example that underscores the importance of the diversity in positions in predicting the effects of variants. Although the alternating amino acid W was not observed at position 93 of the protein Q9UQ88, its position was highly diverse, with 16 amino acids observed. These two examples show that MSA alone did not provide sufficient information for predicting the effects of variants.

The third and fourth examples showed pathogenic variations. The third example involved a substitution from P to L, and although L was not observed in the MSA, all three tools failed to correctly classify this variation. Although the third example was also mislabeled by MSAboost, the fourth example, which corresponded to a substitution from M to T at the first position of the protein with UniProt ID O60500, was labeled as pathogenic by both PHACTboost and MSAboost. There are two reasons for this. First, we used an MSA-based approach to masking (see the “Methods” section for details) for unaligned regions and applied a special rule for the first position. If there was any amino acid other than M at the first position, we masked it (i.e., assigned a gap character instead of the given amino acid). Thus, the frequency of T of the MSAboost algorithm was zero because we used the masked MSA. Second, as explained in the “Methods” section, we added a set of pathogenic variations to the training set that labeled any substitution at the first position as pathogenic. Thus, MSAboost correctly classified this variation through our adjustments, and this demonstrates the benefit of carefully handling of alignment-related problems.

### PHACTboost outperforms AlphaMissense, EVE, and CPT-1

We now compare PHACTboost with several novel and well-known tools that are not part of the dbNSFP: AlphaMissense (Cheng et al., 2023), EVE (Frazer et al., 2021), and CPT-1 (Jagota et al., 2023). Fig. 3A–C show comparisons between AlphaMissense, EVE, and CPT-1, and PHACTboost over all alternative test sets, respectively. The results were consistent across all comparisons, with PHACTboost delivering better performance on all alternative test sets from the easiest to the hardest variants. We observed the same trend in comparisons of the AUPR as before (see Supplementary Fig. 2). Further details such as the AUROC and AUPR of tools, as well as the number of neutral and pathogenic variants in each comparison are listed in Supplementary Tables 6 for each subset generated. We should note that the scores of EVE were unavailable for 19% of the variations in 3,187 proteins, leading to a reduced number of variations considered for the corresponding analyses.

**Figure 3.**
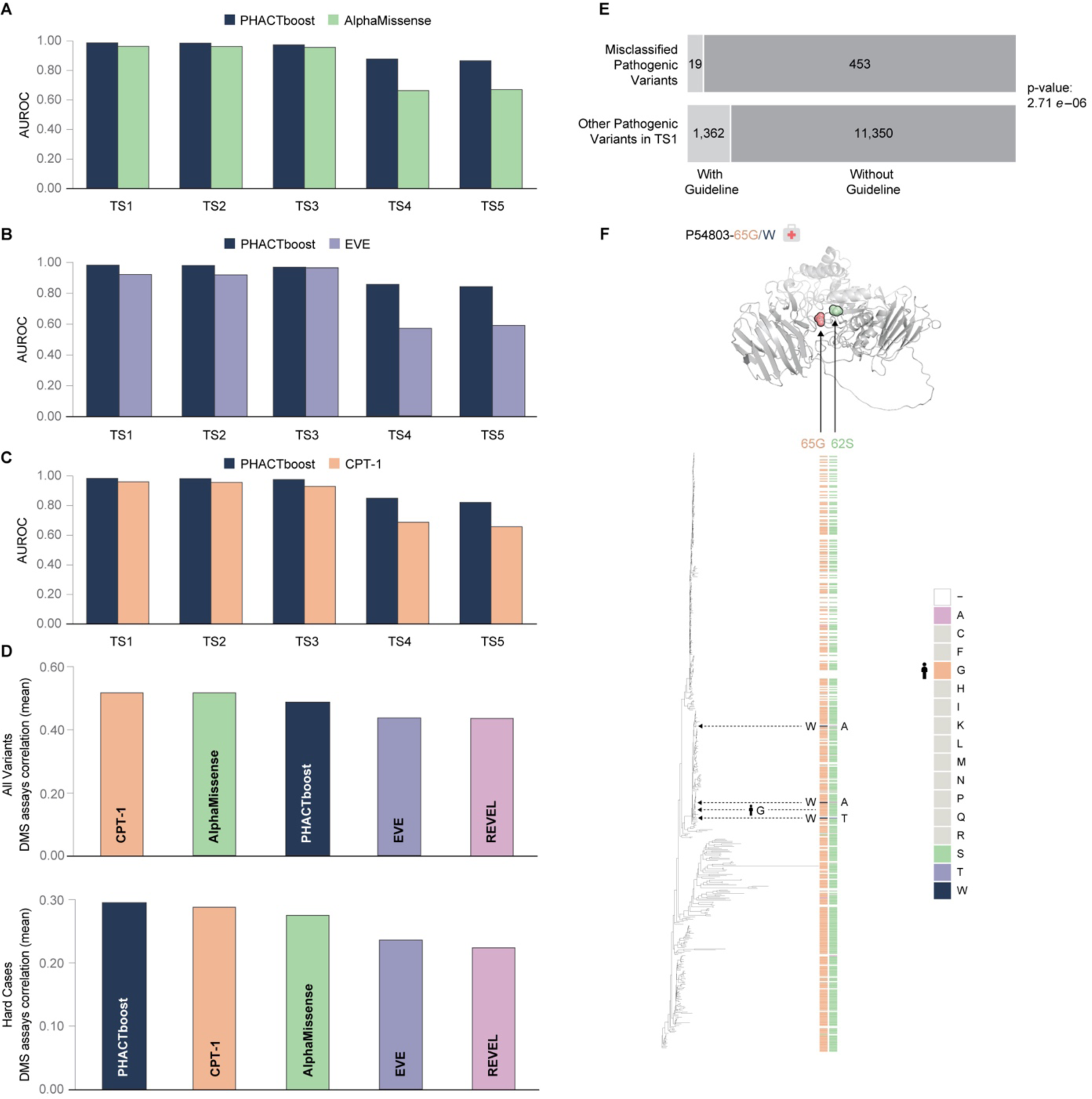
**A.** Comparison of AUROC levels between PHACTboost and AlphaMissense on different alternatives of the test set. **B.** Comparison of AUROC between PHACTboost and EVE on all alternative versions of the test set. **C.** Comparison of AUROC between PHACTboost and CPT-1 on different alternatives of the test set. **D.** Comparison of the average Spearman correlation between PHACTboost and the results of DMS, with the average Spearman correlation between the other tools and DMS data over all variants and hard cases. **E.** Table displaying the number of pathogenic variants submitted to ClinVar and categorized into two groups: those following an ACMG guideline (With Guideline/Without Guideline), and their corresponding accuracy of labeling by PHACTboost (Mislabeled/Others). **F.** An example of a pathogenic variant that was incorrectly classified as neutral by PHACTboost and PHACT due to the omission of the coevolution signal.

### Correlation with DMS experiments

We now determine how well the scores of PHACTboost were correlated with the results of DMS experiments. To test the difference in performance between them, we chose eight genes, and plotted the average Spearman correlation coefficient between all tools and their DMS scores, as in Ref. (Gao et al., 2023). The genes used in the DMS comparisons were as follows: amyloid beta (Seuma et al., 2021), ADRB2 (Jones et al., 2020), MSH2 (Jia et al., 2021), PTEN (Jia et al., 2021; Mighell et al., 2018), BRCA1 (Starita et al., 2015), TP53 (Giacomelli et al., 2018), SYUA (Newberry et al., 2020), and VKOR1 (Chiasson et al., 2020). These genes were chosen because they have been frequent used in recent studies involving predictions of the effects of variants and the comparison of tools based on DMS data (Frazer et al., 2021; Gao et al., 2023).

Figure 3D illustrates that PHACTboost recorded comparable performance to that of AlphaMissense, CPT-1, EVE, and REVEL across all reported variants. It had a correlation coefficient of 0.485, while those of AlphaMissense, CPT-1, EVE, and REVEL were 0.515, 0.514, 0.436, and 0.433, respectively. To assess performance on a subset of challenging cases, similar to the construction of TS4, we curated a “hard cases” subset comprising variants with conflicting pathogenicity predictions from the mentioned four tools. Over this subset, PHACTboost showed a slightly better correlation, as depicted in the second part of Figure 3D. The mean Spearman correlation coefficient was 0.296 for PHACTboost, whereas it was 0.289, 0.275, 0.237, and 0.227 for CPT-1, AlphaMissense, EVE, and REVEL, respectively.

### Analysis of variants misclassified by PHACTboost

In this final subsection, we delve into the specifics of variants that PHACTboost failed to correctly classify. We examined both pathogenic and neutral variations misclassified by PHACTboost with scores close to one and zero, respectively. We identified a noteworthy pattern among them: about 96% of the pathogenic variants mislabeled by PHACTboost had not been submitted to ClinVar following ACMG guidelines. The ACMG offers a structured framework for assessing the clinical significance of genetic mutations or variations, particularly in the context of genetic testing and diagnosing genetic disorders. The number of pathogenic variants that were not covered by these guidelines in our test set was nearly nine times larger than in the other sets, and this disparity grew to 25 times in pathogenic variants that PHACTboost had failed to identify correctly. The table in Fig. 3E provides comprehensive information regarding the total number of pathogenic variants in our test set. These variants were classified into two groups: those following a specific guideline, and those not adhering to any guideline. Moreover, the table indicates whether these variants were accurately identified by PHACTboost by distinguishing between correctly labeled and mislabeled cases (PHACTboost score < 0.5). We used a chi-square test to confirm that there was a significant difference (p-value = 2.71*e* − 06) between variants without ACMG guidelines in TS1 and those mislabeled by PHACTboost. This observation underscores the challenge posed to PHACTboost by pathogenic variations that lack a reliable framework for classification, and ultimately impact the overall reliability of the labels.

In addition to guideline-related information, we observed that the presence of coevolution between protein positions could be a misleading signal of pathogenicity and caused PHACT and PHACTboost to mislabel variants as neutral. While we independently observed alternating amino acids in close proximity to the query protein on the phylogenetic tree, their deleterious effects could be compensated for in these species due to mutations occurring at other positions. This phenomenon aligns with the concept of “compensated pathogenic deviations” described in Ref. (Kondrashov et al., 2002). They claimed that roughly 10% of the differences between a protein in a non-human species and its corresponding human ortholog consist of compensated pathogenic deviations (CPDs). These CPDs arise from amino acid substitutions that, if found at the same site in humans, would be considered pathogenic. We provide an illustrative case from our test set in Fig. 3F. The protein with UniProt ID P54803 exhibited a G-to-W alteration at position 65, which was labeled by ClinVar as pathogenic. However, as illustrated in Fig. 3F, the presence of W at this position was recurrent in a closely related species to humans, suggesting a neutral signal. On the contrary, positions 62 and 65 exhibited a coevolutionary relationship and were in proximity, with a distance of 10.989 based on the 3D structure obtained from AlphaFold and shown in Fig. 3F. Whenever W appeared at position 65, an amino acid different from the reference amino acid S was obtained at position 62. This suggests that these species with W at position 65 may tolerate it owing to an additional mutation at position 62. The failure of PHACTboost to identify these variants resulted from its omission of coevolutionary information.

## Discussion

We present PHACTboost, a gradient-boosted tree-based classifier that uses PHACT scores and features derived from phylogenetic trees, to classify missense variants into pathogenic and neutral classes. We evaluated PHACTboost on a reliable dataset of variations from ClinVar (Landrum et al. 2016) and gnomAD (Karczewski et al. 2020). We also used variants from the human variations dataset (HSV) to eliminate variations with conflicting interpretations. The HSV contains variations obtained from a diverse set of sources, such as the 1000 Genomes Project (Siva, 2008), ExAC (Karczewski et al., 2017), and COSMIC (Forbes et al., 2006). We used the HSV to correctly identify the variations that had been classified differently in the literature with respect to the labels assigned to them. Our comparative analyses showed that PHACTboost outperformed all meta-predictors reported in the dbNSFP. Most of the tools reported in the dbNSFP do not use phylogenetic information in their predictions, because of which they ignore the common evolutionary histories as well as the independence or dependence of alterations in amino acids. As is depicted in Fig. 1C, the use of the MSA alone is not sufficient to obtain the highest accuracy in terms of predictions of pathogenicity, even when using a machine learning-based approach.

Conversely, we observed that VEST4 and PHACTboost delivered similar results on one of the test sets. However, PHACTboost had a clear advantage over all prevalent approaches reported in the dbNSFP for variants that were difficult to predict. That is an important outcome because ClinVar is biased in favor of variants that can be predicted based on evolutionary conservation (Capriotti & Fariselli, 2022). Moreover, because the results of tools to predict the effects of variants were considered to be supporting evidence with respect to ACMG guidelines, the construction of a set of variants that were difficult to interpret represents an important conclusion regarding the performance of these tools. Fig. 2 shows that PHACTboost outperformed all tools, with a significant difference in performance that highlights the importance of phylogeny-based evolutionary information. Fig. 1C shows that PHACT scores constituted the most important feature in predicting the effects of the variants, followed by tree-based features. The results of this study lead us to conclude that phylogenetic trees are an important input for tools for predicting pathogenicity because they are constructed by considering the unique evolutionary history of the genes in question.

To further verify the performance of PHACTboost, we compared it with other popular tools reported in the literature. The results of our comparisons, which were conducted over different test sets, demonstrated that PHACTboost outperformed AlphaMissense, EVE, and CPT-1. The superior performance of PHACTboost, a gradient based tree algorithm, over deep learning-based approaches is noteworthy. It demonstrates that we can enhance the accuracy of tools to predict the effects of variants without using a significant amount of computational resources, which are demanded by deep learning-based methods. Moreover, PHACTboost exhibited better or comparable performance to that of AlphaMissense, EVE, and REVEL, which are three well-known and popular approaches that deliver superior performance to prevalent computational tools, in comparisons in terms of the Spearman correlation with the DMS data.

It was challenging to compare PHACTboost with prevalent tools because we did not have access to the training data used for some of them. This can result in models being tested on data that they might have encountered during their training and can yield optimistic evaluations of their performance. To address this issue, we meticulously segregated variations in ClinVar based on the dates of their submission. However, the lack of analogous information for benign variants from databases like gnomAD restricted the use of this approach. Moreover, currently available tools might have had access to such additional databases as the HGMD and Humsavar, which are not publicly accessible. This might have led to optimistic evaluations of these tools without access to their training data.

In addition to comparing our tool with currently available ones, we investigated the variants that had been mislabeled by PHACTboost. We observed that a significant fraction of the pathogenic variations that were mislabeled by PHACTboost had been submitted to ClinVar without considering any ACMG guideline. As stated in the “Results” section of this study, the ratio of the pathogenic variants that were classified based on guidelines to other pathogenic variants was eight in entire test set and was 25 for pathogenic variants that were predicted as neutral by PHACTboost with a significant score. We also observed that some of the variations that were mislabeled as neutral by PHACT and PHACTboost had in fact been influenced by coevolution. A majority of current approaches overlook coevolution and treat each position as an independent entity. While a select few, such as EVmutation (Hopf et al., 2017) and GEMME (Laine et al., 2019), do incorporate patterns of coevolution in predicting the functional consequences of missense mutations, they rely on the frequency and conservation of the MSAs to identify the coevolving positions, thereby neglecting evolutionary history. In our future research, we aim to incorporate the coevolution of positions into both PHACT and PHACTboost.

We also identified a crucial problem with the ClinVar databases during our research that highlights a potential bias in the evaluations of computational tools. Many prevalent studies have relied on using two summary files of ClinVar variants before and after a specified date, and have considered variations exclusive to the recent file as post-date additions to ClinVar. However, we found that some variants that were present in older versions of ClinVar had been omitted in subsequent versions and reintroduced later. This inconsistency might have resulted in the inadvertent incorporation of variants that had been specified after a certain date into the test set such that they might have been present in the training sets of the other tools. To counter this, we referred to the submitted summary file that detailed the dates of submission of certain variants, and analyzed 53 ClinVar data files that included all variants reported from 2015 to 2020. Variants within this timeframe were used as our training set. Many recent studies appear to have missed this intricacy, and this has possibly led to an overestimation of the performance of the relevant tools. This revelation underscores the need for transparency and methodological rigor when appraising and contrasting computational tools to predict the effects of variants.

While PHACTboost delivered comparable average correlation with the DMS data as AlphaMissense, EVE, REVEL, and CPT-1 in our analysis, we must acknowledge the ambiguity associated with the absolute contribution of DMS assays in predicting the effects of variants. While DMS experiments serve as an important piece of evidence, they are just one of the many sources that are consulted to gauge the potential effects of variants on protein function. Incorporating DMS data into computational models, as demonstrated in our study, offers valuable experimental validation. Nevertheless, it is crucial to interpret the results within the context of their inherent limitations and potential biases.

Another avenue for future research involves considering paralog-related information to determine the weights of ancestral nodes in PHACT scoring. Gene duplication plays an important role in the acquisition of new functions (Long et al., 2003). Following gene duplication, one of the paralogs may accumulate more mutations and functionally diverge from the other (Ohno, 1970). Integrating functionally divergent sequences into the MSA and the phylogenetic tree may help identify substitutions that are intolerable in functionally divergent lineages but tolerable in the original lineage. Our algorithm does not explicitly consider this difference at present, but we partially cover the divergence of one copy by assigning a lower weight to its corresponding node by using the weight function in PHACT. In future work, we plan to incorporate the process of gene duplication into PHACT and add related features to PHACTboost. This will reduce the importance of the second copy through the identification and conservation of the ancestral function of the relevant node.

## Methods

### Data Collection and Processing

#### BLAST & multiple sequence alignment

To identify homologues for each query sequence, we employed PSI-BLAST (Position-Specific Iterated BLAST) (Altschul et al., 1997) against a non-redundant database of 14.010.480 proteins, derived from the reference proteomes in the UniProtKB/Swiss-Prot Knowledgebase (UniProt, 2021). We perform two iterations of PSI-BLAST with 5000 maximum target sequences. Due to computational limitations associated with building phylogenetic trees, we capped the number of hits to a maximum of 1000 sequences. Additionally, to ensure sequence quality, we imposed a minimum identity threshold of 30% and an E-value threshold of 0.00001. The sequences are aligned using MAFFT FFTNS (Katoh & Standley, 2013), and the MSA is trimmed with the trimAl tool gappyout method (Capella-Gutierrez et al., 2009).

#### Maximum-likelihood phylogenetic tree

The resultant MSA is used to generate a maximum-likelihood phylogenetic tree with the FastTree (Price et al., 2010). tool, with LG model and leaving the other parameters at their default settings. Using FastTree, we constructed phylogenetic trees for 20,191 proteins.

#### MSA masking

Though MAFFT is a trusted tool for MSA construction, we still observed unaligned regions, especially in areas in gapped blocks. To reduce alignment problems, we employed an MSA masking approach. This strategy focuses on two scenarios: i) identifying incompatible regions, wherein amino acids are present only once at a position, and over half of its immediate neighboring positions (5 preceding and 5 following) are inconsistent with the rest of the segment. Ii) identifying incompatible regions before and after gap breaks.

For the latter, we check if the starting and ending amino acids of the block are aligned correctly. If either one aligns, indicating it is not the sole amino acid in that position, we refrain from masking any part of the block. Conversely, if this criterion isn’t fulfilled, we mask the entire block, substituting by masked amino acids with gaps. This MSA masking step is essential, as the quality of MSA significantly impacts the accuracy of ASR (Vialle et al., 2018). Therefore, we apply the masking scheme before performing ASR.

#### Ancestral reconstruction

Positions with a ‘gap’ character in the query sequence are removed from the initial MSA. The refined MSA is then employed for ASR using IQTREE with the R4 model, where an equal substitution rate is applied to all amino acids, while other parameters are set to their default settings.

#### Construction of the variant set

We assembled a dataset comprising verified neutral and pathogenic variations from ClinVar and gnomAD. We also incorporated data from the *Homo sapiens* variations (HSV) database (UniProt, 2023), which encompasses human variants from databases such as ExAC, TOPMed, to eliminate the variants with conflicting interpretations in ClinVar. To map genomic coordinates from ClinVar and gnomAD to protein positions, we developed a tool, Map2Prot. This pipeline takes the GRCh38 genomic coordinates of ClinVar and gnomAD variations and provides the corresponding protein position and the type of the variant (synonymous, non-synonymous, early stop codon, etc.). The details of this pipeline can be found in the Supplementary Materials.

ClinVar variant_summary.txt file dated 23.02.2023 includes 1,922,312 GRCh38 coordinates, with 1,397,809 mappable to our protein positions. From ClinVar and gnomAD v3, we obtained 934,009 and 3,580,039 non-synonymous variations, respectively. To refine the dataset, we considered the benign variants labeled as “benign” and/or “likely benign,” and the pathogenic variants labeled as “pathogenic” and/or “likely pathogenic”. We removed variants with labels other than “benign”, “likely benign,” “pathogenic” and “likely pathogenic”. To further eliminate variants with conflicting interpretations from the literature, we utilize HSV, which includes variations from different databases. In the subsequent step, we combined single nucleotide-level variations corresponding to the same protein into a single variation. Although rare, we also remove the same protein variant with different interpretations from the variant set.

To ensure the quality of the pathogenic and neutral class labels, we applied allele frequency (AF) thresholds to the remaining variants. Variants were classified as neutral if their AF from ClinVar or gnomAD was greater than or equal to 0.01. Pathogenic variations were retained only if their AF was reported in gnomAD and was ≤ 0.005. Notably, the primary source of pathogenic variations is ClinVar, meaning that we include only clinically verified pathogenic variants in our analyses.

To conduct a fair comparison with other variant effect predictors, the test set exclusively comprises variants from the ClinVar January 2020 release. This approach aims to exclude variants present in the training sets of either individual or meta-predictors from our test set as much as possible. Meta-predictors incorporate outputs of other pathogenicity prediction tools, while standalone predictors as tools do not. Importantly, we realized that deciding on the submission date of the variations by retrieving them from ClinVar before and after a certain date is not sufficient to identify the submission date of the variations accurately. To address this, we used the submission summary file provided by ClinVar, along with all variant summary files from 2015 to 2019 (consisting of 59 files). By checking whether the submission date given in submission_summary.txt is before 2020 or if a variant exists in one of the 59 variant_summary.txt files, we determined whether to include it in the training set. Variants not meeting these criteria were assigned to the test set. Regarding the benign variants from gnomAD, since the submission date is not available, we randomly divided them into train and test sets to achieve a balanced ratio of neutral to pathogenic variations.

Furthermore, we introduced 10,000 pathogenic missense mutations that involve variations from the reference amino acid (RefAA) at the first position of proteins. This step aimed to teach the model to classify variations to any other amino acid as pathogenic for the first position. Additionally, we randomly selected 10,000 RefAA-RefAA substitutions and labeled them as neutral. These adjustments were made to instruct the model to identify reference amino acid substitutions as neutral. In total, we added all these 20,000 variants to the training set to enhance the model’s learning capabilities.

To ensure quality of the evaluation set, we excluded from the test set that had a ClinVar review status of zero stars. The final version of the training set included 13,062 proteins and 70,937 variants. After this exclusion, the test contains 27,310 variants. This test set (referred to as TS1) comprises 14,116 neutral and 13,194 pathogenic variants, associated with 8,423 distinct proteins.

As the training and test folds of most existing tools are not accessible, to minimize variant overlap with their training and test data, we separated ClinVar variants into distinct test sets based solely on their submission dates. However, while the variant set is not overlapping, this splitting strategy results in having proteins present in both the training and test sets. Such variants could be easier to predict if the protein-specific features are sufficient to predict the variant class label. Therefore, we created additional test sets. TS2 includes variants located on protein positions not observed in PHACTboost’s training set and TS3 comprises variants from proteins that are not present in the training data.

To assess the performance of PHACTboost and other tools on hardest cases, we constructed TS4 and TS5 subsets, which encompass variants where 40 tools sourced from dbNSFP disagreed. Specifically, variants on which at least 25% of these tools predicted class label (pathogenic or neutral) are discordant, with no consideration to the actual class label. TS4 dataset includes all variants satisfying this criterion, whereas TS5 is composed of mixed proteins included in TS4 that exhibit both neutral and pathogenic variants. In that regard, TS5 includes the hardest variants to predict for PHACTboost.

To further assess PHACTboost on the test variants that do not cause optimistically successful predictive performances, we created subsets of TS1 based on the pathogenic-to-neutral ratio in variants of a protein observed in the training set, namely TS1_Pure, TS1_Mixed, TS1_0.1-0.9, TS1_0.2-0.8, TS1_0.3-0.7, TS1_0.4-0.6. TS1_Pure includes proteins with only pathogenic or neutral variants in the training set, whereas TS1_Mixed includes proteins with both pathogenic and neutral variants in the training set. TS1_0.1-0.9, TS1_0.2-0.8, TS1_0.3-0.7, and TS1_0.4-0.6 include proteins with a pathogenic-to-neutral ratio in variants observed in the training set between 0.1 and 0.9, 0.2 and 0.8, 0.3 and 0.7, 0.4 and 0.6, respectively. The details of these subsets, including the number of proteins and neutral and pathogenic variants, are given in Supplementary Table 7.

#### Workflow engine and high-performance computing (HPC)

Conducting a comprehensive, reproducible, and scalable data analysis for multiple proteins with various parameters on an Hight Performance Computing (HPC) center is not straightforward (Wratten et al., 2021). Each task (job) exhibits distinct characteristics. For example, generating maximum-likelihood phylogenetic trees with FastTree tool is a CPU-intensive process, whereas tasks like Blast and constructing MSA are memory-intensive. Most tasks can be completed within hours, however generating phylogenetic trees may take up to 8 hours. The HPC environment is complex, consisting of hundreds of servers operating simultaneously, and occasional failures are not uncommon. Managing a large number of jobs in such an environment can be challenging. Therefore, it is required to employ a workflow tool that defines the analysis through a set of rules and generates output files from input files. This approach not only ensures scalability but also enhances reproducibility, allowing other researchers to obtain the same results effortlessly. All tools and software used during the analyses, input files, the computational facilities can be specified in a text file. This enables easy deployment of the analysis environment without additional effort.

To satisfy all these requirements, we used a Snakemake workflow (Mölder et al., 2021) with a conda package manager due to its human-readable, Python-based language, portability, integration with a conda package manager, automatic deployment, and ability to specific software dependencies. In this study, 20,191 proteins with ten consecutive tasks mean 201,910 independent jobs were executed using Snakemake workflow and almost 1.6M CPU hours resource used in a HPC cluster ss PHACTboost takes average 10 hours with 8 cores to compute tolerance score per protein.

We retrieved 1,806 eukaryotic proteomes from the UniProt reference proteomes database (Release 2021_03). Our pipeline was applied to a total of 20,588 proteins belonging to Homo sapiens. During the analysis, we excluded 334 of proteins due to their limited number of PSI-BLAST hits, falling below the threshold of 10 hits.

### Experimental Setting

#### Machine learning algorithm

We used a gradient boosting-based machine learning algorithm, LightGBM (G. Ke et al., 2017) to train our prediction model. We implemented PHACTboost in R using the LightGBM’s R package version 3.2.2.

#### Hyper-parameter tuning for LightGBM

LightGBM has many hyper-parameters affecting the performance of the algorithm. We selected the following hyper-parameters for LightGBM from the given ranges in Supplementary Table 8 using a 4-fold cross-validation strategy on the training dataset. Since applying a grid search on the whole hyper-parameter set simultaneously would be computationally infeasible, we followed a sequential tuning approach to pick the model hyper-parameters. We first initialized each hyper-parameter and then, using the defined ranges in Supplementary Table 8, we tuned and updated each hyperparameter sequentially by picking the value with the best mean cross-validation AUROC until no other hyperparameters were left. We set the number of boosting iterations and the early stopping rounds parameters as 5000 and 250, respectively.

#### Performance measures

We used area under receiver operating characteristic (ROC) curve (AUROC) and area under precision-recall curve (AUPR) to evaluate the predictive performance of pathogenicity prediction algorithms. In our analyses, true positive and true negative correspond to positive (i.e., pathogenic) and negative (i.e., neutral) labels, respectively.

### PHACTboost Input Features

We trained PHACTboost on several feature groups as PHACT and PHACT-related scores, MSA, phylogenetic tree, ASR-based features, and amino acid classes. In Supplementary Table 2, we list the details of all features included in the model. There are 427 features in total used as input for the PHACTboost.

## Data and Code Availability

The all data generated in this study, all benchmark analysis scripts and source codes for PHACTboost are available at https://github.com/CompGenomeLab/PHACTboost.

## Acknowledgements

This work was supported by the Health Institutes of Turkey (TUSEB) (Project no: 4587 to O.A.) and EMBO Installation Grant (Project no: 4163 to O.A.) funded by the Scientific and Technological Research Council of Turkey (TÜBİTAK). Most of the numerical calculations reported in this paper were performed at the High Performance and Grid Computing Center (TRUBA resources) of TÜBITAK. The TOSUN cluster at Sabanci University was also used for computational analyses. We also want to thank Nehircan Özdemir for his art illustration of the PHACTboost approach.

## SUPPLEMENTARY MATERIAL

### Map2Prot – Mapping GRCh38 genomic coordinates to the protein positions

To align genomic coordinates of variations with corresponding protein positions, we developed a pipeline known as Map2Prot. This pipeline systematically processes genetic annotation data in GTF format and genome sequence data to extract coding sequence (CDS) information from gene transcripts. We used the “Homo_sapiens.GRCh38.dna.primary_assembly.fa.gz” Ensembl’s release 106 files as our reference genome. This dataset was sourced from the Ensembl database’s FTP site, accessible at: https://ftp.ensembl.org/pub/release-106/fasta/homo_sapiens/dna/. The gene annotation data, formatted in GTF, was acquired from the “Homo_sapiens.gtf” file, available in release 106 on the Ensembl database’s FTP site at: https://ftp.ensembl.org/pub/release-106/gtf/homo_sapiens/. This dataset includes each gene’s chromosome, strand, transcripts, exon details, and start-stop coordinates, encompassing only protein-coding genes, translated processed pseudogenes, and translated unprocessed pseudogenes.

The extraction of CDS sequences was facilitated by employing the “samtools” version 1.9’s (Danecek et al., 2021).

**Supplementary Figure 1:**
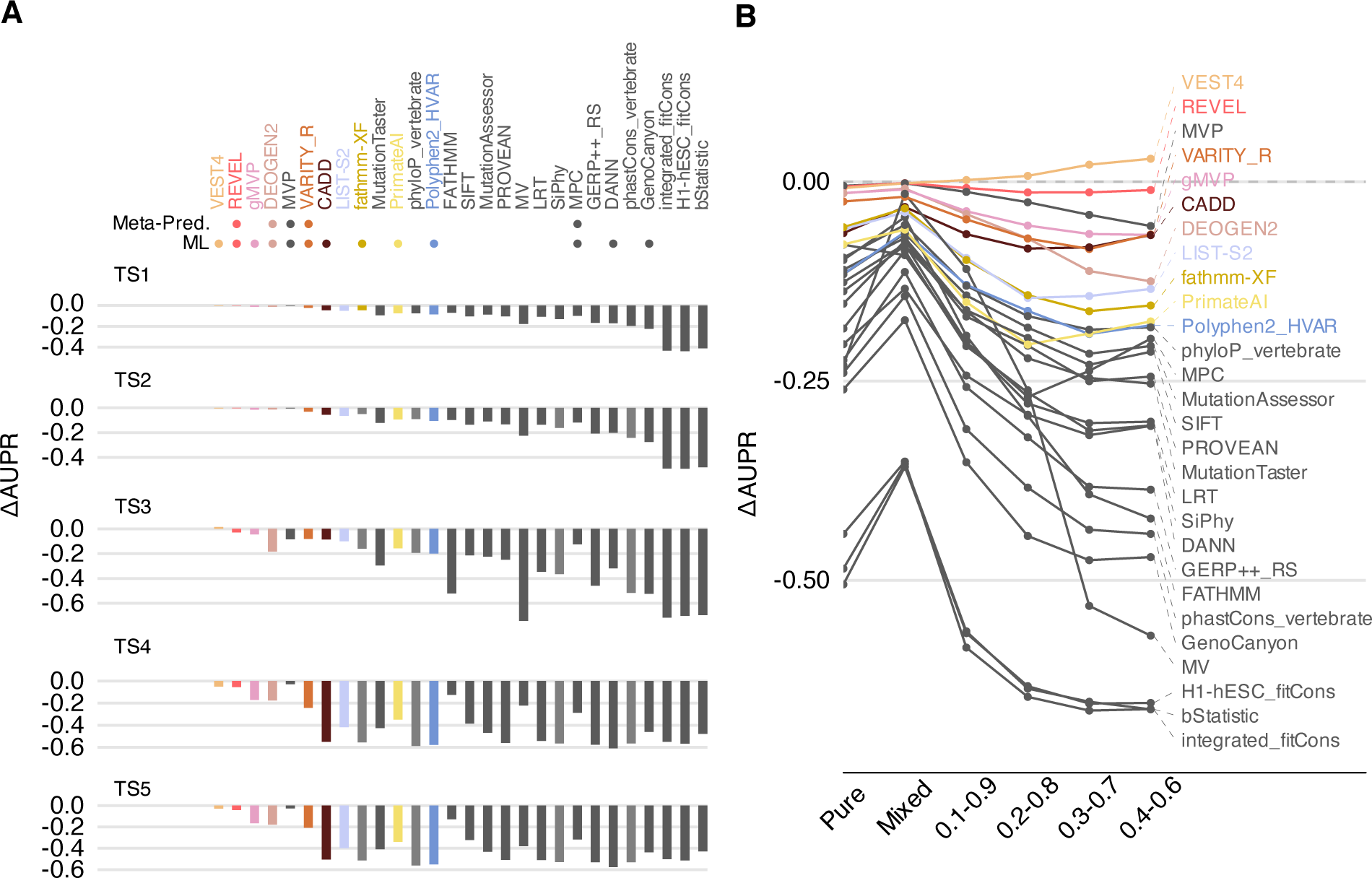
AUPR comparisons of PHACTboost against tools reported in dbNSFP and PMV. A-E. Comparisons across TS1, TS2, TS3, TS4 and TS5, respectively. F. Comparisons across TS1, TS1_Pure, TS1_Mixed, TS1_0.1-0.9, TS1_0.2-0.8, TS1_0.3-0.7, and TS1_0.4-0.6. G. Comparisons across TS4, TS4_Pure, and TS4_Mixed. ΔAUPR corresponds to the difference between AUPR levels of tools and PHACTboost, negative values correspond better predictive performance of PHACTboost.

**Supplementary Figure 2.**
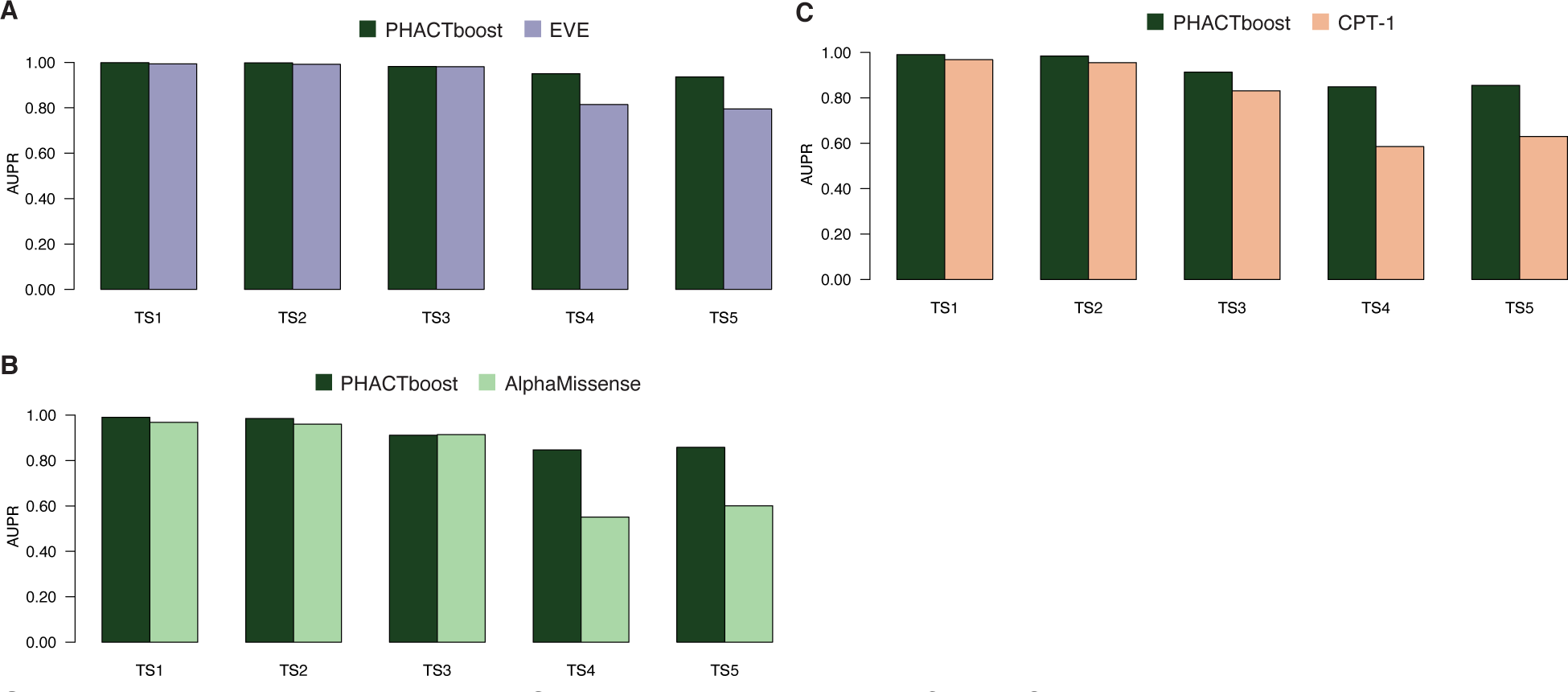
**A, B, C.** AUPR comparisons of PHACTboost and EVE, AlphaMissense and CPT-1, respectively.

